# The Hippo pathway effector Taz is required for cell fate specification and fertilization in zebrafish

**DOI:** 10.1101/304626

**Authors:** Chaitanya Dingare, Alina Niedzwetzki, Petra A Klemmt, Svenja Godbersen, Ricardo Fuentes, Mary C. Mullins, Virginie Lecaudey

## Abstract

In the last decade, Hippo signaling has emerged as a critical pathway integrating extrinsic and intrinsic mechanical cues to regulate cell proliferation and survival, tissue morphology and organ size *in vivo*. Despite its essential role in organogenesis, surprisingly much less is known about how it connects biomechanical signals to control of cell fate and cell size during development. Here we unravel a novel and unexpected role of the Hippo pathway effector Taz (*wwtr1*) in the control of cell size and cell fate specification. In teleosts, fertilization occurs through a specific structure at the animal pole, called the micropyle. This opening in the chorion is formed during oogenesis by a specialized somatic follicle cell, the micropylar cell (MC). The MC has a peculiar shape and is much larger than its neighboring follicle cells but the mechanisms underlying its specification and cell shape acquisition are not known. Here we show that Taz is essential for the specification of the MC and subsequent micropyle formation in zebrafish. We identify Taz as the first *bona fide* MC marker and show that Taz is specifically and strongly enriched in the MC precursor before the cell can be identified morphologically. Altogether, our genetic data and molecular characterization of the MC lead us to propose that Taz is a key regulator of the MC fate activated by physical cues emanating from the oocyte to initiate the MC morphogenetic program. We describe here for the first time the mechanism underlying the specification of the MC fate.

## Introduction

The molecular and cellular mechanisms driving cell fate specification and cell differentiation have largely engaged developmental biologists in the last decades. Acquisition of a specific cell fate results from a combination of biochemical and physical cues originating from the cell itself or from its direct or more distant environment. One of the longstanding questions in the field is how these cues are interpreted in a combinatorial manner by cells to adopt a cell fate different from their neighbors. To address this question, we used the zebrafish ovarian follicular epithelium as a model system.

Fertilization is a key event in the life cycle of all sexually reproducing organisms, but is achieved differently in different species. In teleost fish, a sperm penetrates the oocyte through a narrow canal called the ‘micropyle’ that traverses the thick vitelline membrane. In zebrafish, the micropyle consists of a vestibule followed by a cone-shaped canal opening up onto the egg surface. Since the inner aperture of the micropyle is only slightly larger than the size of a sperm head, the micropyle also represents a mechanical barrier against polyspermy (Hart and Donovan, 1983). The micropyle is made by a specialized follicle cell, the micropylar cell (MC), generated during the process of oogenesis and around which vitelline material is deposited (Hart and Donovan, 1983; Kobayashi and Yamamoto, 1985; Nakashima and Iwamatsu, 1989; Selman et al., 1993).

Oogenesis in zebrafish can be divided into five stages (Selman et al., 1993) (Fig. 1). During stage I, individual oocytes become surrounded by follicle cells (FCs), and components of the vitelline membrane begin to accumulate between the oocyte and the FCs. At that stage, an aggregate of RNA, proteins and organelles called the Balbiani body forms on the future vegetal side of the oocyte and determines the oocyte animal-vegetal (AV) axis (Marlow and Mullins, 2008). During Stage II, FCs proliferate and adopt a cuboidal shape to cover the growing oocyte. Both FCs and the oocyte extend microvilli that contact each other via cell-cell junctions. During stage III, yolk precursor protein and lipids accumulate in the oocyte, and the MC becomes morphologically distinguishable from other FCs. At the end of stage III, the oocyte has almost reached its mature size. During stage IV, the germinal vesicle migrates to the animal pole and the oocyte and FCs retract their microvilli prior to ovulation. Finally during stage V, the mature egg, free of FCs, is released, and can be fertilized by the sperm through the micropyle (Selman et al., 1993).

**Figure 1.**
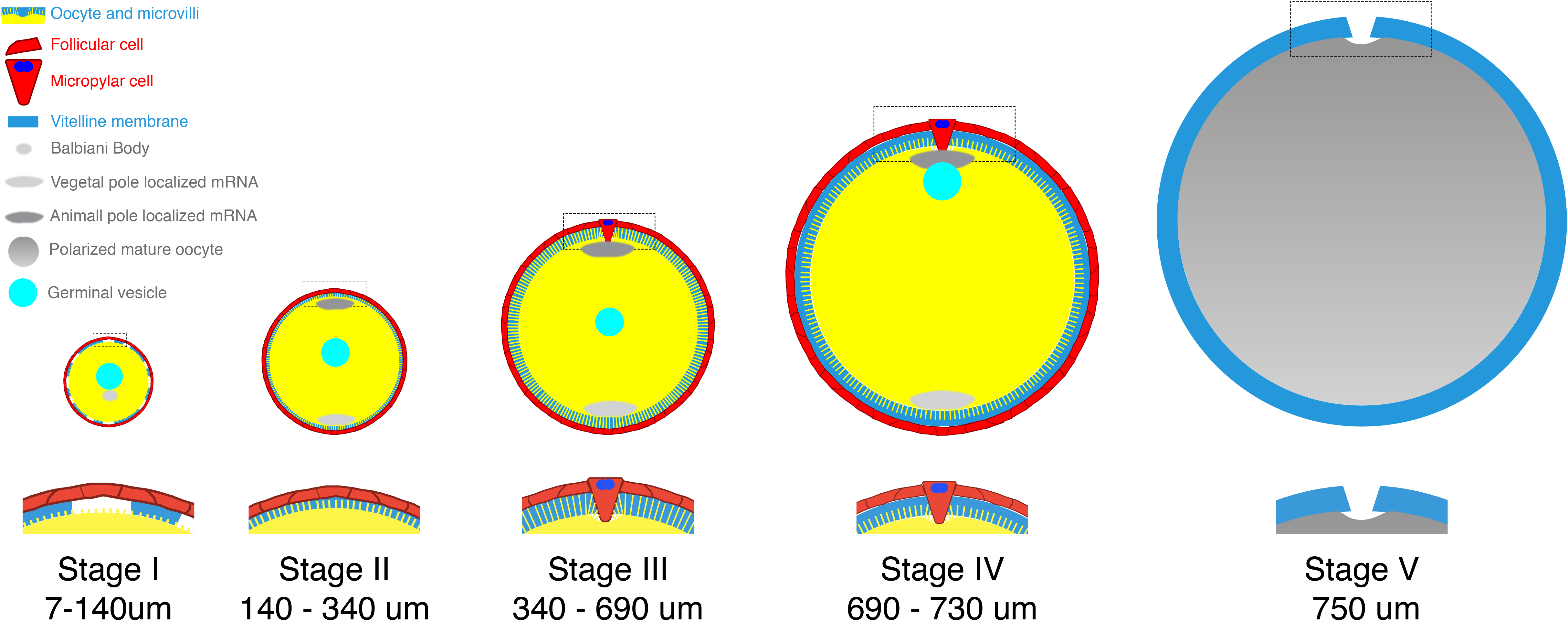
Scheme recapitulating the five different stages of zebrafish oogenesis (based on Selman et al., 1993) Late stage I: Follicular cells (FC, red) surrounding the single-cell oocyte (yellow), and components of the vitelline membrane (blue) accumulate in between. The Balbiani body (grey) forms and establishes the oocyte animal-vegetal (AV) axis. Stage II: Microvilli start to grow at the surface of the oocyte. Different RNAs are specifically localized at the vegetal or animal (light and dark grey, respectively). Stage III: The MC becomes morphologically distinguishable from other FCs. Stage IV: The germinal vesicle (turquoise) migrates to the animal pole. The oocyte microvilli retract prior to ovulation. Stage V: Mature, FC-free egg surrounded by the vitelline membrane, which is perforated by the micropyle at the animal pole. Close-up of the animal pole are displayed under each follicle scheme. Diameter sizes of the oocytes are indicated.

How can a modified FC perforate the vitelline membrane? Although little is known on the detailed structure of the MC in zebrafish, EM analyses in Medaka and in the Chum salmon have provided the first insights into how the MC morphology can determine the micropyle architecture (Kobayashi and Yamamoto, 1985; Nakashima and Iwamatsu, 1989). From stage III of oogenesis onward, the MC becomes much larger and adopts a typical mushroom-shape that easily distinguishes it from other FCs. The proximal part of the “mushroom” (closest to the oocyte) forms a cytoplasmic extension, which contains intertwined bundles of microtubules and tonofilaments, and remains in contact with the oocyte cell surface, also known as the oolemma. The cytoplasmic extension is thought to either prevent accumulation of vitelline material and/or to apply mechanical pressure on the growing vitelline membrane. Finally, the MC shrinks at the end of oogenesis and withdraws from the micropylar canal thus leaving a hole in the vitelline membrane (Fig. 1, Stage V) (Kobayashi and Yamamoto, 1985; Nakashima and Iwamatsu, 1989). Despite this detailed ultra-structural description, no biological marker for the MC has been identified so far. Furthermore, almost nothing is known about the genetic and molecular mechanisms leading to MC specification.

The MC invariably forms at the animal pole of the oocyte. Oocytes become polarized along the AV axis at the end of stage I of oogenesis and the Balbiani body plays a crucial role in this process by determining the vegetal pole of the oocyte. In zebrafish *bucky ball (buc)* mutant oocytes, the Balbiani body fails to form causing oocyte polarity loss, with the oocyte displaying animal pole identity radially expanded and lacking the vegetal pole (Marlow and Mullins, 2008; Bontems et al., 2009). As a result, multiple micropyles form around the oocyte, demonstrating the importance of oocyte polarity establishment to spacially restrict the MC (Marlow and Mullins, 2008). Thus, the abnormalities observed in the *buc* mutant phenotype suggest that the wild-type oocyte signals to the FC layer to position and induce the specification of the MC. Despite this first observation, the mechanisms underlying the selection and the differentiation of the MC remain completely unknown.

The Hippo signaling pathway was discovered less than 15 years ago in *Drosophila* in mutants showing giant organs (Huang et al., 2005, reviewed in Dong et al., 2007; Pan, 2007). The core Hippo kinase cascade leads to the phosphorylation of the pathway effector Yorkie and to its sequestration and/or degradation in the cytoplasm (Huang et al., 2005). In contrast, when the core kinase cascade is inactive, Yorkie can translocate into the nucleus and, together with the transcription factor Scalloped, activate the transcription of genes promoting cell proliferation, growth or survival. This core Hippo kinase cascade (also referred to as “canonical”) is fully conserved in vertebrates with the 2 paralogs Yap (Yes-Associated Protein) and Taz (Transcriptional co-Activator with PDZ-binding motif), two homologs of Yorkie, being the Hippo pathway effectors (Zhao et al., 2007, reviewed in Varelas, 2014; Zhao et al., 2008; Zhao et al., 2011). Remarkably, Yap and Taz function also as oncogenes and their expression and nuclear localization are increased in several human cancers (reviewed in Moroishi et al., 2015; Zygulska et al., 2017).

Recently, many other proteins associated with cell junctions and/or apico-basal polarity have been proposed to limit Yap and Taz function independently of the core kinase cascade and thus constituting the “non-canonical” Hippo pathway (reviewed in Lv et al., 2017; Sasaki, 2015; Stamenkovic and Yu, 2010). Molecular components of the canonical and the non-canonical cascades interact with each other to control the localization and the activity of Yap/Taz and hence to limit proliferation and organ size. In addition to their role in limiting organ size, Yap and Taz are involved in other cellular processes including cell fate specification. YAP is, for example, essential for the specification of trophectoderm versus inner cell mass fate in the mammalian blastocyst downstream of both cell polarity and cell adhesion (Hirate et al., 2013; Leung and Zernicka-Goetz, 2013; Manzanares and Rodriguez, 2013; Nishioka et al., 2009). In addition, YAP and TAZ have been involved in controlling fate specification in human embryonic and pluripotent stem cells (Beyer et al., 2013; Musah et al., 2014), zebrafish retinal pigment epithelial cells (Miesfeld et al., 2015) and various other tissues (Judson et al., 2012; Yimlamai et al., 2014; Zhang et al., 2011).

Here we investigate the role of the Hippo pathway effector Taz in zebrafish oogenesis and report an unexpected and novel function for Taz in the specification of a single, peculiar FC, the MC. We show that the Hippo pathway effector Taz encoded by the *wwtr1* gene (WW domain-containing transcription regulator protein 1) is a key regulator of MC specification in zebrafish. We further show that *wwtr1/taz* mutant females lay eggs that cannot be fertilized because they lack a micropyle. While *wwtr1* mutant oocytes seemed to progress normally throughout oogenesis, we found that the MC failed to differentiate within the follicular cell layer surrounding the oocyte. We show that Taz is strongly enriched in the future MC before morphological changes, making Taz the first *bona fide* marker of this peculiar, somatic cell type. Our findings suggest that Taz is required to specify the MC during early oogenesis. By using cell junctions and cytoskeletal molecular markers, we molecularly characterized the MC for the first time. These results open the door for gene functions discovery in micropyle biogenesis. We propose a working model linking oocyte polarity, Taz function and MC fate determination.

## Results

### *wwtr1* mutant females are infertile

To address the function of Taz during embryonic development of the zebrafish, we used a reverse genetic approach, the Transcription Activator-Like Effector Nuclease (TALEN), to target the 1^st^ exon of the *wwtr1* gene. We generated several mutant alleles including *wwtr1^fu55^* (thereafter *wwtr1^-/-^* or *wwtr1* mutant refers to *wwtr1^fu55/fu55^*). The induced genomic lesion lead to a 8 nucleotides deletion and a 1 nucleotide insertion, creating a predicted premature stop codon after the first 29 amino acids (Fig. 2A-C). The zygotic mutant embryos did not show any obvious morphological phenotype and grew to adults that were indistinguishable from their siblings (Fig. 2D, E).

**Figure 2.**
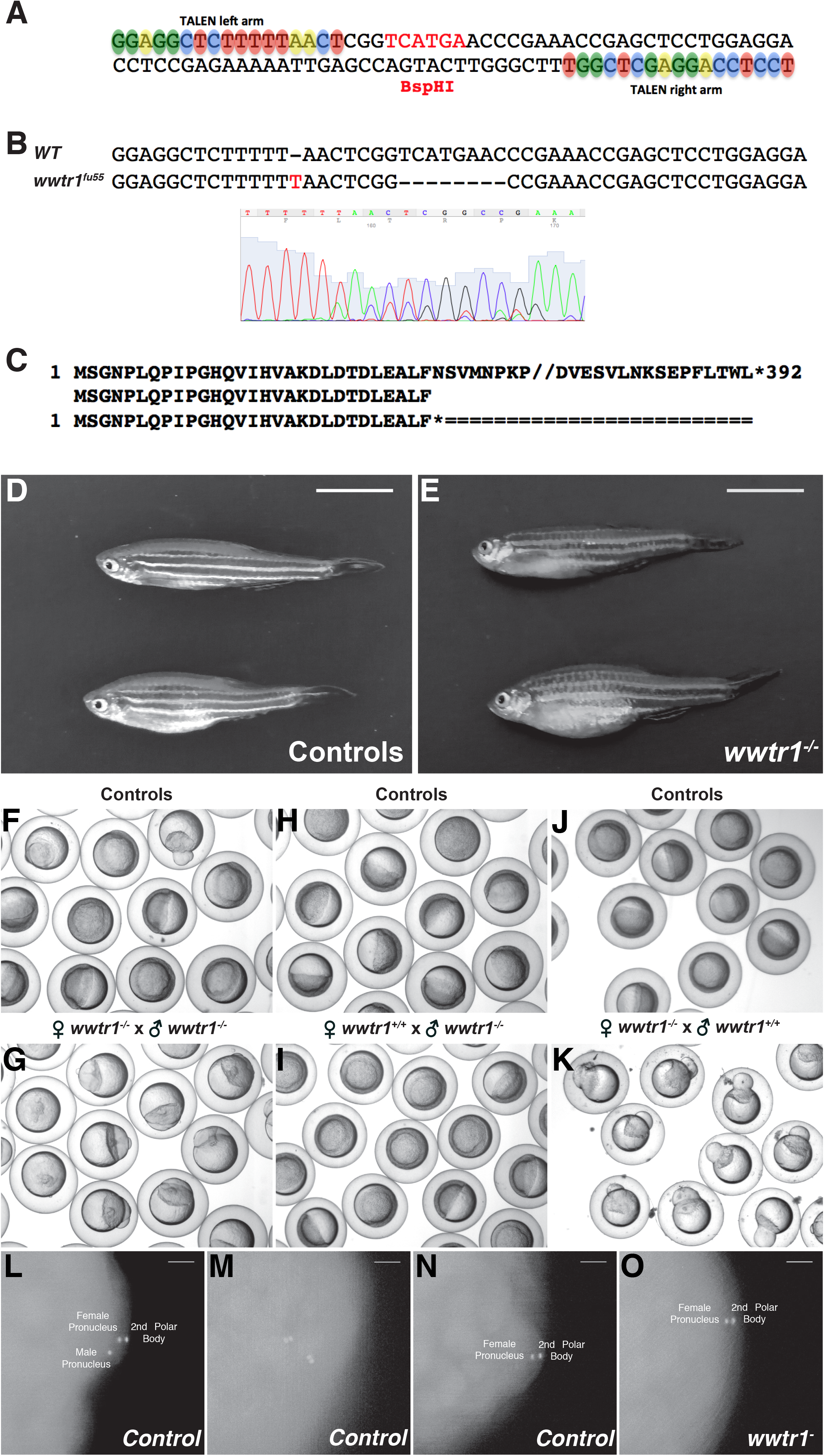
*wwtr1* mutant females are infertile. (A) TALEN target sequence within the *wwtr1* locus (exon1). Left and right TAL repeats are highlighted and color-coded. The BspHI restriction site within the spacer region, used for screening and genotyping, is marked in red. (B, C) Molecular characterization of the *wwtr1^fu55^* allele. The *fu55* allele has one inserted and 8 deleted base pairs within or in the vicinity of the spacer region (B) leading to the generation of a premature STOP codon after 29AA (C). (D, E) Homozygous adult mutant fish (E) do not show any obvious morphological phenotype as compared to WT or heterozygous siblings (D). (F-K) Eggs or shield stage embryos from a *wwtr1^-/-^* incross (G, N=3), an outcross of *wwtr1^-/-^* males to WT females (I, N=3), or an outcross of *wwtr1^-/-^* females to WT males (K, N=3), compared to sibling controls (F, H and J). Eggs laid by *wwtr1^-/-^* females systematically fail to undergo cleavage independent of the male genotype (n > 100). (L-O) WT (L-N, N=2, n=40) or *wwtr1^-^* mutant (O, N=2, n=40) eggs were fertilized *in vitro* with WT sperm, fixed after 10 minutes and stained with DAPI to label the nuclei. Most of the control eggs showed 3 DAPI positive structures corresponding to the male and female pronuclei, and to the second polar body (L, 60%). Occasionally, 4 or only 2 DAPI positive structures were visible due to polyspermy (M, 11%) or absence of fertilization (N, 29%), respectively. In contrast, *wwtr1* mutant eggs systematically showed only 2 DAPI positive structures corresponding to the female pronucleus and to the 2^nd^ polar body (O, 100%) showing that these eggs were not being fertilized. N is the number of experiments, n is the number of eggs. Scale bars are: 1 cm in D-E, 100 μm in L-O

To test a potential maternal or paternal contribution of *wwtr1*, homozygous mutant females were crossed to homozygous mutant males while wild-type siblings were incrossed as controls (Fig. 2F,G). All eggs from the *wwtr1^-/-^* mutant incrosses failed to undergo cleavage and appeared as unfertilized eggs (Fig. 2G). To further test whether this fully-penetrant phenotype resulted from the mother or the father genetic contribution, *wwtr1^-/-^* males were crossed to WT females (Fig. 2I, related control in 2H) and *wwtr1^-/-^* females were crossed to WT males (Fig. 2K, related control in 2J). While eggs from crosses between *wwtr1^-/-^* males and WT females underwent cleavage normally (Fig. 2I), eggs from crosses between *wwtr1^-/-^* females and WT males did not undergo cleavage and appeared as unfertilized eggs (Fig. 2K). To make sure that the observed phenotype was due to the mutation in *wwtr1*, we performed a complementation experiment by combining in trans the *wwtr1^fu55^* allele and the previously published *wwtr1^mw49^* alleles (Miesfeld et al., 2015). While eggs from *wwtr1^mw49/+^* females crossed to WT males developed normally (Supp. Fig.1A-C), none of the eggs laid by *wwtr1^mw49/fu55^* females underwent cleavage (Supp. Fig.1D-E). Altogether, these results suggest that the maternal absence of Taz affects the eggs’ competence to be fertilized.

We next tested whether *wwtr1* mutant eggs could be fertilized. To this end, we performed *in vitro* fertilization (IVF) assays of *wwtr1* or control egss with WT sperm. Eggs were fixed after 10 minutes and stained with DAPI. In 60% of the control eggs, 3 DAPI-positive dots could be identified corresponding to the male and female pronucleus and DNA of the 2^nd^ polar body (Fig. 2L). Four and two DAPI-positive dots were observed in 11% and 29% of the cases, respectively, corresponding to either polyspermy events often reported in IVF assays (Fig. 2M) or unfertilized eggs (Fig. 2N). In contrast, in 100% of the *wwtr1* mutant eggs, only two nuclei were detected, corresponding to the female pro-nucleus and the 2^nd^ polar body DNA (Fig. 2O), identical to unfertilized eggs (compared to Fig 2N). This result indicated that eggs laid by homozygous *wwtr1* mutant females cannot be fertilized.

### *wwtr1* mutant eggs lack a functional Micropyle

To determine the cause of this infertility, we dissected out ovaries from *wwtr1* mutant females and control siblings (henceforth control refers to either WT or *wwtr1^+/-^* siblings). The overall morphology of the ovaries was indistinguishable in control siblings (Fig. 3A) and in *wwtr1* mutant females (Fig. 3B), excluding the possibility that the infertility issue might be caused by a defective oogenesis. As mentioned above, the sperm penetrates the oocyte through the micropyle in teleost fishes. To test if the infertility of *wwtr1* mutant females could be due to a defect in the formation of the micropyle, mature eggs were squeezed from gravid females and activated in embryo medium. While the micropyle could be easily identified as a small dark spot on the surface of the inflated chorion in control (arrow in Fig. 3C, C’), such a structure was never visualized in *wwtr1* mutant eggs (Fig. 3D, D’). To further confirm the lack of micropyle, activated eggs were stained with coomassie brilliant blue (CBB), which binds to glycoproteins enriched in the micropylar structure (Yanagimachi et al., 2013). A darkly stained spot on the outer surface of the chorion, corresponding to the micropyle, was clearly visible in activated control eggs (arrow in 3E, E’) but not mutant eggs (Fig. 3F, F’). These results show that eggs from *wwtr1^-/-^* females fail to be fertilized because the chorion lacks a micropyle.

**Figure 3.**
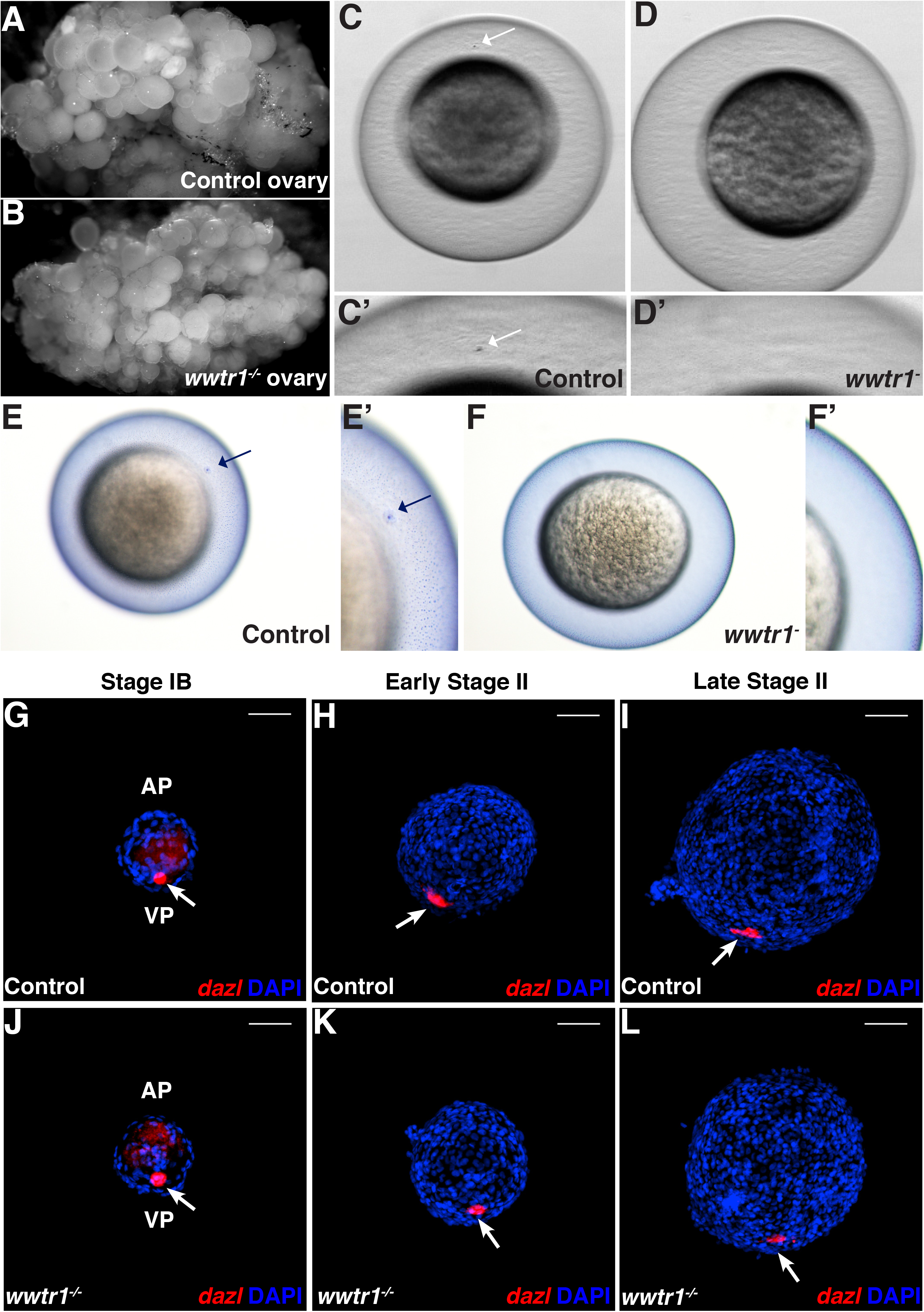
Eggs from *wwtr1* mutant females do not form a micropyle but show proper animal-vegetal polarity establishment. (A-B) Whole ovaries form siblings (A, n=2 ovaries) and *wwtr1^-/-^* females (B, n=2 ovaries) showing comparable morphology. (C-D’) Activated control egg with the easily identifiable micropyle at the surface of the chorion (C, C’ white arrow, N=3, n=20). A comparable structure cannot be observed in the *wwtr1* mutant egg (D, N=3, n=20). C’ and D’ images are close-up images of C and D, respectively. (E-F’) Coomassie Brilliant Blue staining of the micropyle in an activated control (E, n=10) or *wwtr1* mutant egg (F, n=8). While in control, a darkly stained micropyle can be observed (E, E’), no such spot could be observed in the mutant (F, F’). E’ and F’ are close-up of E and F, respectively. (G-L) Whole-mount *in situ* hybridization with the vegetal pole and Balbiani body marker *dazl* in control (G-I, white arrow, n=24) and *wwtr1* mutant (K-L, white arrow, n=22) oocytes at the indicated stage of oogenesis, showing that mutant oocytes establish a normal AV polarity and form a Balbiani body (J-L) like controls (G-I). FC nuclei are stained with DAPI. n is the number of ovaries, eggs or follicles. AP, animal pole; VP, vegetal pole. Scale bar is: 50 μm in G-L

### *wwtr1* mutant oocytes exhibit normal animal-vegetal polarity

In the zebrafish *bucky ball (buc)* mutant, in which AV polarity fails to be established, supernumerary micropyles form due to radial animal identity, leading to polyspermy (Marlow and Mullins, 2008). To test whether the absence of a micropyle in *wwtr1* mutant eggs could be due to a loss of animal identity leading to a loss of the animal pole-localized micropyle, we investigated AV polarity formation in *wwtr1* mutant oocytes. We performed whole-mount *in situ* hybridization (WISH) on dissected follicles at different stages to detect the localization of *dazl* mRNA, one of the first transcripts to localize asymmetrically in the oocyte (Elkouby et al., 2016; Maegawa et al., 1999). The *dazl* mRNA localizes to the Balbiani Body as soon as it forms in stage IB of oogenesis and later resides at the vegetal pole of the oocyte. The *dazl* localization pattern in mutant oocytes at different stages was indistinguishable from that in control oocytes (compare Fig. 3J-L to G-I, white arrows), indicating that the Balbiani body forms properly in *wwtr1^-/-^* oocytes. In addition, we examined the localization of *vg1* mRNA which localizes to the animal pole in stage III oocytes when the MC is fully formed (Bally-Cuif et al., 1998; Marlow and Mullins, 2008). In *buc* mutants, *vg1* mRNA is found in multiple domains beneath each of the multiple micropyles, suggesting that the localization of *vg1* and the micropyle at the animal pole is interdependent (Marlow and Mullins, 2008). Surprisingly, *vg1* localization was not affected in *wwtr1^-/-^* oocytes and remained localized in one spot at the animal pole like in control oocytes (supp. Figure 2A-B). Altogether, these results indicate that the absence of micropyle in *wwtr1* mutant eggs is not due to a disruption of oocyte AV polarity, and that conversely the absence of the micropyle does not seem to affect oocyte polarity.

### The micropylar cell fails to differentiate in the absence of Taz activity

The micropyle is generated by the MC, a specialized somatic FC, that differentiates at the animal pole of the FC layer. We thus tested whether defective MC formation could cause the absence of micropyle in *wwtr1* mutant eggs. To test this possibility, we investigated MC development during oogenesis in control and *wwtr1^-/-^* females. In wild-type ovaries cryosections, labeled with Rhodamine-phalloidin to detect F-actin, MCs were fairly easy to identify in late stage III and older oocytes based on their typical mushroom-shape (Fig. 4A, green arrow). Interestingly, on the oocyte side, a superficial indentation associated with F-actin accumulation was evident at and around the contact point with the MC (Fig. 4A, red arrow). In *wwtr1^-/-^* ovaries, in contrast, no large, mushroom-shaped cell could be identified within the follicular cell layer. Yet, a large indentation could be observed at the oocyte surface associated with actin accumulation similar to that observed in control oocytes (Fig. 4B, red arrow). In this case, however, instead of being interrupted by the MC like in control, the microvilli that cover the oocyte formed a continuum within this superficial indentation (Fig. 4A-B, blue arrows). This result suggests that the MC is absent from the FC layer in *wwtr1^-/-^* follicles.

**Figure 4.**
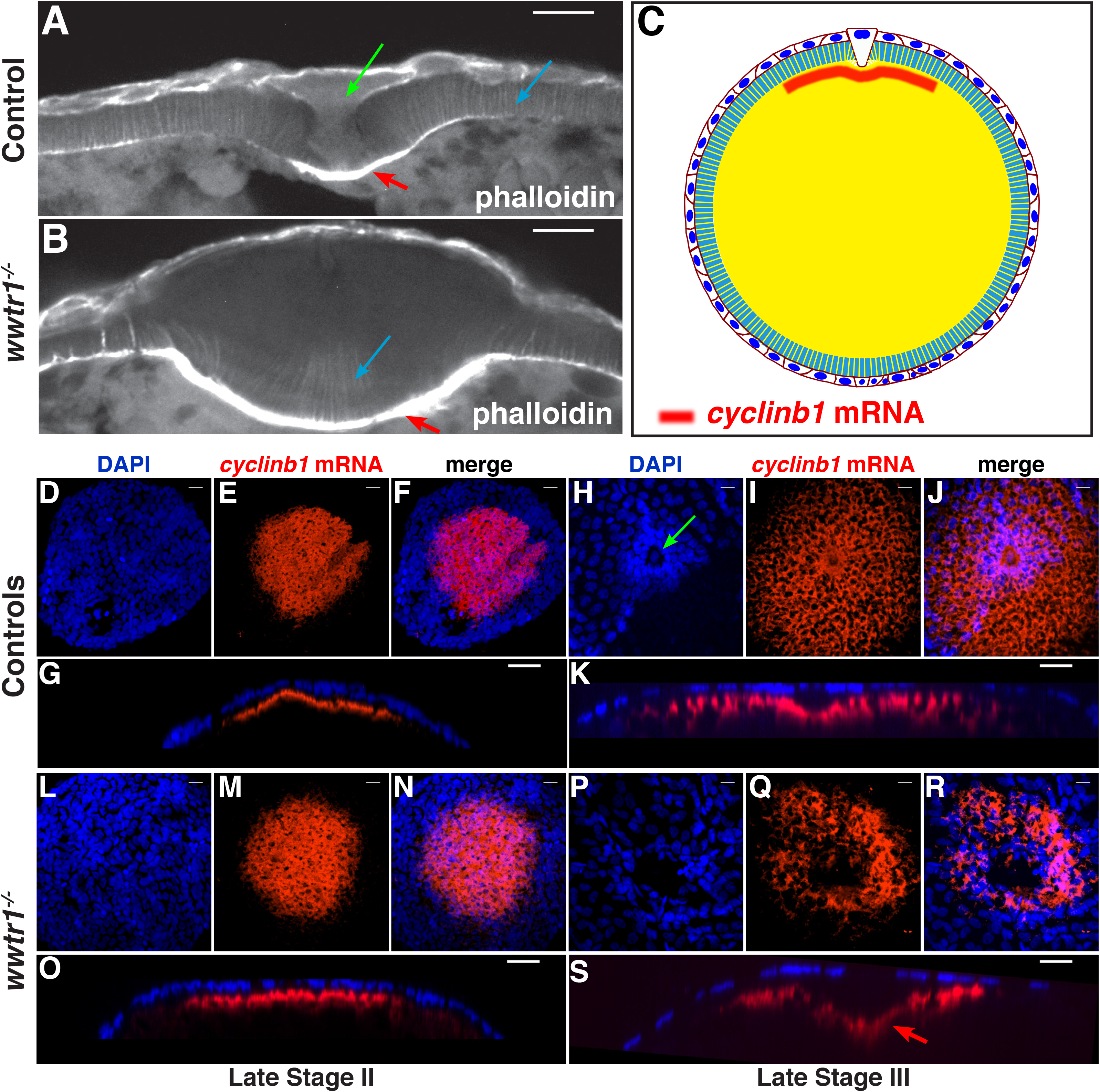
*wwtr1* mutant oocytes lack the micropylar cell. (A-B) Confocal images projection of control (A) and *wwtr1^-/-^* (B) cryosectioned ovaries stained with Rhodamine-Phalloidin. The typical mushroom-shaped MC is easily identifiable in the control ovaries (A, green arrow), while such typical structure is not present in the mutant oocyte (B). Blue arrows point to the microvilli formed between the FC layer and the oocyte. Red Arrows indicate the surface indentation and actin enrichment at the animal pole surface in both control and *wwtr1* mutant oocytes. (C) Schematic of a stage III oocyte showing *ccnb1* mRNA localization at the animal pole and the MC in the FC layer surrounding the oocyte. (D-S) Confocal images projection of controls (D-K, n=31) or *wwtr1^-/-^* (L-S, n=40) oocytes stained by *in situ* hybridization to detect *ccnb1* mRNA at the animal pole (red), combined with DAPI nuclear staining (blue) at the indicated oocyte stages. At the stage II oocyte, the FC layer is homogeneously organized at the animal pole in both controls (D, F, G) and *wwtr1* mutant (L, N, O). During stage III, control FCs adopt a typical concentric arrangement around the centrally located MC (green arrow in H). In contrast *wwtr1* FCs get disorganized at this stage at the animal pole and the MC cannot be identified (P-S). (G, K, O, S) are X-Z orthogonal views of corresponding confocal stacks (F, J, N, R). The red arrow in panel S shows the pronounced indentation at the animal pole of the oocyte in the absence of a MC in *wwtr1^-/-^* oocytes. n is the number of follicles. Scale bars are: 10 μm for A-B, D-F, H-J, L-N and P-R, and 20μm for G, K, O and S.

To confirm this result, we combined WISH with a probe against the animal pole marker *ccnbl (cyclinbl)* and DAPI staining to label the FC nuclei (Fig. 4C). At late stage II of oogenesis, the earliest stage at which *ccnb1* mRNA localized to the animal pole, the FC layer was homogeneously organized at the animal pole both in controls (Fig. 4D-G) as well as in *wwtr1^-/-^* oocytes (Fig. 4L-O). In the stage III oocyte, control FCs at the animal pole assumed a typical concentric arrangement around the centrally located MC (Fig. 4H-K, green arrow). In contrast, *wwtr1^-/-^* FCs showed a disorganized animal pole arrangement at this stage and no MC could be identified (Fig. 4P-S).

Instead, a large indentation was present in the animal pole cortex in *wwtr1^-/-^* oocytes (Fig. 4S, red arrow) similar to that observed in the cryosections (Fig. 4B). This large indentation was in the center of the *ccnb1* positive cytoplasmic domain confirming its localization at the animal pole, where the MC should normally differentiate. These results show that Taz activity is required for the formation of the MC within the follicular cell layer.

### Taz is the first *bona fide* marker of the micropylar cell

To understand how Taz induces the MC fate and micropyle formation, we sought to determine whether Taz is expressed in the oocyte or in the FCs. We first performed RT-PCR at different developmental stages including mature oocytes and embryos before and after the maternal-zygotic transition (Supp. Fig. 3). *wwtr1* mRNA was neither present in mature oocytes nor at the 256-cell stage, and started to be detected at 60% epiboly. *wwtr1* mRNA is therefore not maternally provided, suggesting that it may be expressed in the follicular cell layer. To test this hypothesis, we stained stage III oocytes with a commercially available anti-Taz antibody, combined with β-catenin and DAPI staining to label cell membranes and nuclei, respectively. Strikingly, Taz protein appeared highly enriched in one cell at the surface of the FC layer (Fig. 5A). This cell was easily identifiable as the MC based on the regular organization of the other FC nuclei around it (Fig. 5B). In addition, β-catenin, as expected, sharply delineated the membranes of all FCs that were in the same XY plane, except the MC (Fig. 5C-D). An orthogonal projection indeed showed that the Taz positive cell is much bigger as compared to the other FCs (Fig. 5E), in agreement with the large size and mushroom-shape morphology of the MC (Fig. 4A). To confirm that Taz is a *bona fide* marker of the MC, we examined Taz localization in *buc* mutant follicles, which are known to form several MC. At the surface of *buc* oocytes, multiple, distinctly-localized cells were positive for Taz (Fig. 5F-G’). These observations show that Taz is highly enriched in the MC as compared to other follicular cells, identifying Taz as the first molecular marker uniquely distinguishing the MC.

**Figure 5.**
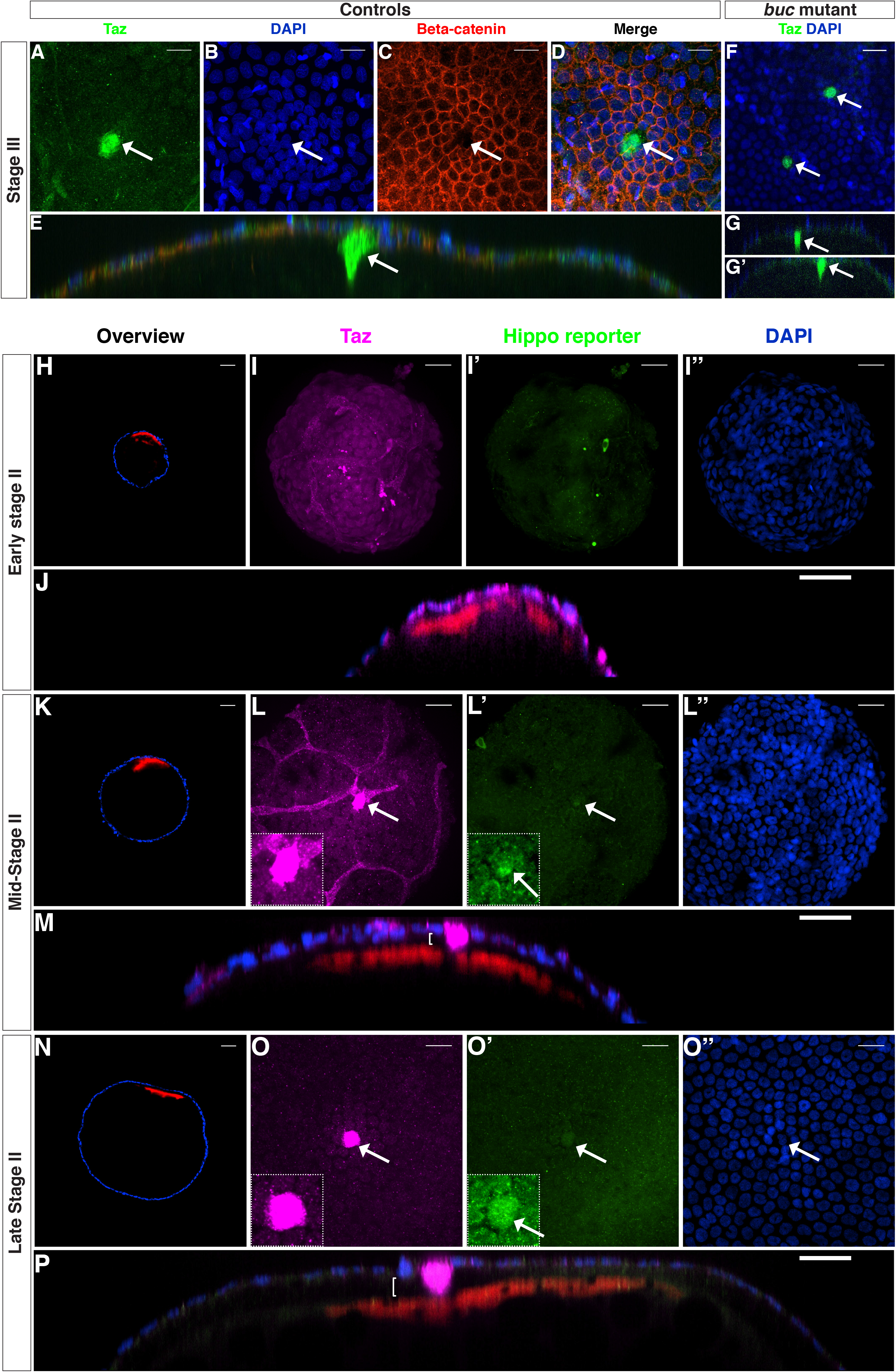
Taz protein is a *bona fide* marker of the MC. (A-E) Confocal images of stage III WT oocytes stained with anti-Taz (A) and anti-β-catenin (C) antibodies, and DAPI (B, n=4). D shows the 3 channels merged, and E is an X-Z orthogonal view of D showing the Taz-positive MC in cross-section. (F-G’’) Stage III *bucky-ball* mutant oocytes stained with the anti-Taz antibody and DAPI (n=4). G and G’ are X-Z orthogonal views of F showing two Taz-positive MCs. White arrows in (A-G’) point to the MCs. (H-P, n=38) are confocal images of whole mount Tg(4xGTIIC:d2egfp)^mw50/0^ oocytes stained with the anti-Taz antibody (magenta), *ccnb1* mRNA (red) and DAPI (blue) at early stage II (H-J), mid-stage II (K-M), and late stage II (N-P). Hippo reporter activity from the Tg(4xGTIIC:d2egfp) transgene is shown in green. XZ sections (J, M, P) show that a gap between the oocyte and the FC layer forms and become larger as development proceeds (white bracket in M and P) except at the level of the MC. n is the number of follicles. Scale bars are: 20 μm for A-D, F, I, J, L, M, O, P and 40 μm for H, K, N.

To gain more insights into the role of Taz in the specification of the MC, we determined how early in oogenesis Taz protein is enriched in the MC, using *ccnb1* mRNA localization to mark the animal pole. Follicles were collected from Tg(*4xGTIIC:d2egfp*)*^mw50^* females expressing the Hippo/TEAD reporter to additionally evaluate Taz activity (Miesfeld and Link, 2014). In early stage II oocyte (Fig. 5H, follicle size 140 μm) Taz protein levels were similar in all FCs and the GFP signal from the TEAD reporter line was very low in the entire FC layer (Fig. 5I-J). From mid-stage II onwards (Fig. 5K, follicle size 210 μm), Taz levels strongly increased and the TEAD reporter exhibited slightly increased activity in the MC precursor as compared to the rest of the FCs (Fig. 5L-M, arrow). In late stage II oocytes (Fig. 5N, follicle size 300 μm), Taz levels remained very high in the MC and the TEAD reporter activity was also clearly distinguishable from the other FCs (Fig. 5O-P, arrows). These results identify Taz as the first marker uniquely distinguishing the MC precursor from the rest of the FCs. Taz is present in the future MC as soon as midstage II, before obvious morphological changes in this cell. These results further suggest that Taz could be one of the very first factors playing a role in the specification of the MC.

### Molecular characterization of the micropylar cell

To better understand the transition from FC to MC and the role of Taz in this process, we further characterized the MC and its contact with the oocyte at the molecular level. A striking difference between the MC and the other FCs is indeed the way each contacts the oocyte surface: while the FCs are thought to be mainly in contact with the oocyte microvilli (Cerdà et al., 1999; Hart and Donovan, 1983; Kessel et al., 1985; Nakashima and Iwamatsu, 1989; Wolenski and Hart, 1987), the MC directly contacts the oolemma at all stages of oogenesis and despite the drastic morphological changes it goes through. We thus aimed at characterizing the MC-oocyte contact point in more detail.

To do so we needed tools to unequivocally mark the MC or the animal pole. For this purpose, we used either the anti-Taz antibody or the *ccnb1* ISH probe to specifically identify the MC and the oocyte animal pole, respectively. However, the anti-Taz antibody could neither be used in the *wwtr1* mutant nor in combination with other antibodies produced in the same animal species (rabbit), and not all antibodies can successfully be used after ISH. Therefore we generated a new transgenic line, Tg(*zp0.5:egfp-zorba*), to express the animal pole-localized Zorba protein fused to eGFP under the control of the oocyte-specific promoter zp0.5 (Bally-Cuif et al., 1998; Onichtchouk et al., 2003). In positive oocytes, eGFP-Zorba localized at the animal pole (Supp Fig. 4). In oocytes from F1 *Tg(zp0.5:eGFP-Zorba)^+/0^* females, however, GFP was never detected suggesting that the transgene was systematically silenced already in the first generation. Oocytes from F0 females were still a very useful tool to simultaneously analyze various junctional and cytoskeletal components.

First, we examined tight junctions using an antibody against Zonula occludens-1 (ZO-1). Tight junctions have been proposed to be present at the contact point between the oocyte and MC in other species (Kobayashi and Yamamoto, 1985). In control oocytes at early stage III, a thick actin accumulation (white arrow in Fig. 6A) was observed in the cortical actin layer just beneath the MC-oocyte contact point but no ZO-1 staining was detected (Fig. 6A’, C, C’, white, blue and yellow brackets in Figure 6 mark the oocyte cortex layer, the oocyte microvilli layer and the FC layer, respectively). In mid-stage III oocytes, the thick actin accumulation persisted at the animal pole of the oocyte beneath the MC (white arrows in Fig. 6B) and ZO-1 appeared as a thin ring, included within the thicker F-actin domain (white open arrowhead in Fig. 6B’, D, D’). In mutant oocytes at the same developmental stage, an indentation was detected at the animal pole of the oocyte as described above (Fig. 4B and 4S), but no MC was associated with this indentation (compare Fig. 6E to 6B). Accordingly, no ZO-1 staining was observed at the oolemma at the animal pole (Fig. 6E’, F, F’). On an animal view, the indentation of the oolemma was clearly visible (Fig. 6F, F’). These results reveal that tight junctions are present between the MC and the oocyte oolemma in control follicles and confirm the absence of MC in *wwtr1^-/-^* follicles.

**Figure 6.**
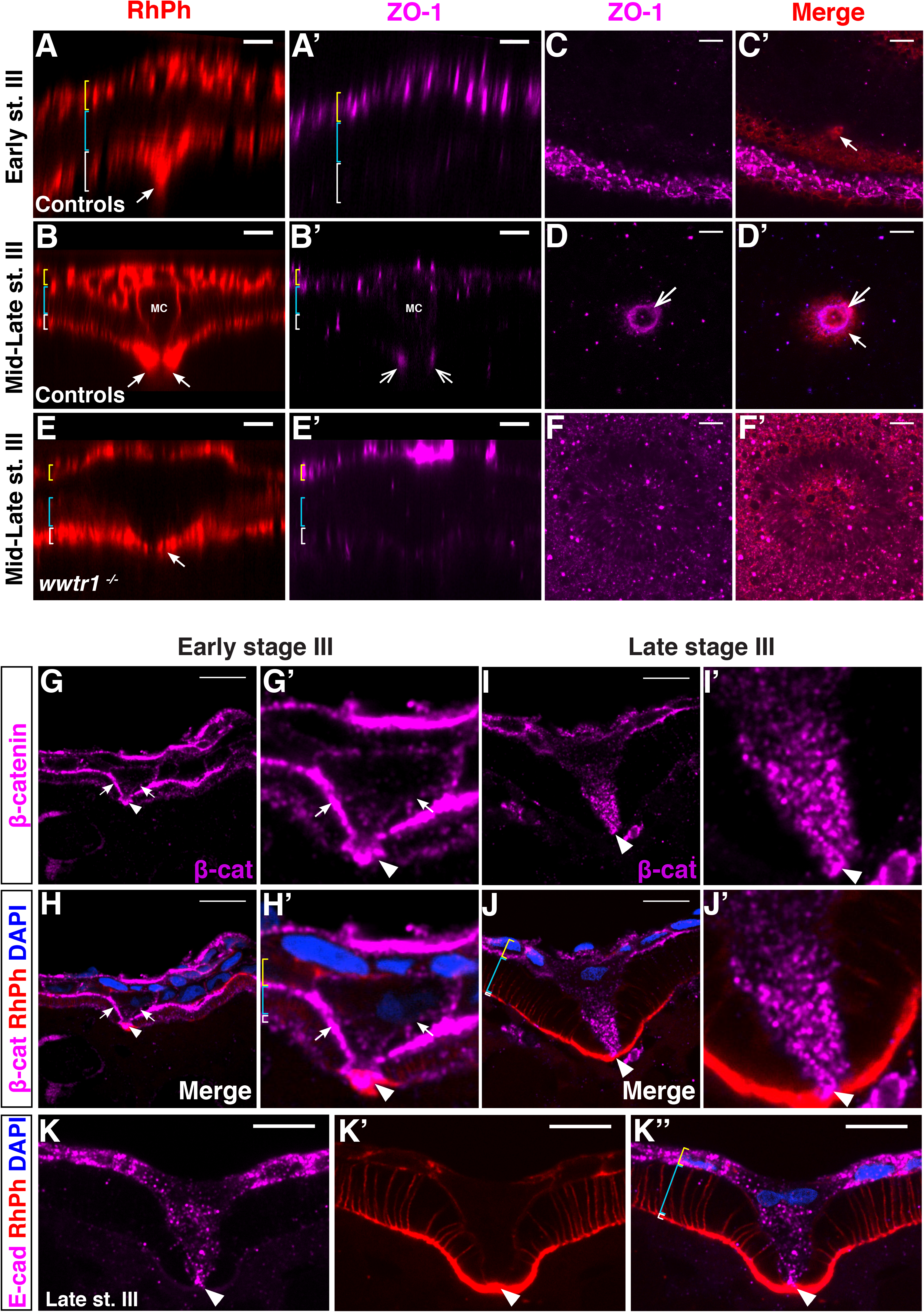
Tight and adherens junctions are established between the micropylar cell and the developing oocyte. (A-F’) Whole-mount controls (A-D’) and *wwtr1^-/-^* (E-F’) follicles stained with Rhodamine Phalloidin to label F-actin (red) and an anti-ZO1 antibody to label tight junctions (Magenta) (N=1, n=7). (A, A’, B, B’, E, E’) are lateral viewed orthogonal sections of the oocyte animal pole with the FC layer on top. (C, C’, D, D’, F, F’) are animal viewed confocal projections (stack size 4 μm) of the oocytes (C and C’ are tilted relative to the animal pole). At both stages, recruitment of actin is visible at the MC-oocyte contact site in controls (solid arrow in A, B, C’, D’). At early stage III oocyte, ZO-1 is present in the FC layer (yellow bracket in A’) but not at the MC-oocyte contact point (A-C’). During mid/late stage III, ZO-1 staining becomes visible as a ring surrounding the MC at the contact point with the oocyte (simple arrow in B’, D, D’). This ZO-1-stained ring coincides with the actin accumulation signal (solid arrow in D’). In *wwtr1^-/-^* follicles (E-F’), actin is present at the animal oocyte cortex but not as strongly enriched as in the control oocytes (Compare white arrows in E and B) and the MC is absent (E). ZO-1 is detected in the FC layer as in controls (yellow bracket in A’, B’, E’) but not at the animal oocyte oolemma (E’, F, F’), in agreement with the absence of MC. Animal views of the follicle show the indentation at the animal pole of the *wwtr1^-/-^*oocyte (F, F’). White, blue and yellow brackets mark the oocyte cortex layer, the oocyte microvilli layer and the FC layer, respectively. (G-J’) Confocal images of cryosectioned ovaries showing β-catenin localization in the MC and the neighboring FCs (n=10). In the early stage III oocyte, β-catenin marks the lateral membrane of the FCs and of the MC (solid arrow), and the contact point between the MC and the oocyte (arrowhead) (G-H’). In the late stage III oocyte, when the MC displays its typical mushroom shape, β-catenin appears disengaged from the lateral membrane except at the contact point between the oocyte and the MC (arrowhead in I-J’). Similar to β-catenin, E-cadherin (n=8) also does not show any lateral membrane staining in the MC but at the contact point between oocyte and the MC at the late stage III (arrowhead in K, K’’). Rhodamine-Phalloidin (red) and DAPI (blue) were used to independently mark for membranes and nuclei. Scale Bar: 10 μm in A-H.

Tight junctions control the permeability of junctions but have poor adhesive properties (Hartsock and Nelson, 2008). We thus used β-catenin and E-cadherin antibodies to determine whether adherens junctions were also present at the MC-oocyte contact point. β-catenin was present at the contact point from the beginning of stage III (arrowhead in Fig. 6G-H’). At this stage, it also localized to the lateral membrane of the MC (arrows in Fig. 6G-H’). In late stage III oocytes, β-catenin was still present at the MC-oocyte contact point (arrowhead in Fig. 6I-J’) but had delocalized from the lateral membrane of the MC and appeared as small aggregates in the cytoplasm (Fig. 6I-J’). E-cadherin was not clearly detected at early stage III, but at late stage III its localization at the MC-oocyte contact point and as small aggregates in the cytoplasm was similar to β-catenin localization (compare Fig. 6I-J’ to K-K’’). These results suggest that β-catenin and E-cadherin function in maintaining the adhesion between the MC and the oolemma in the stage III oocyte, and that during MC differentiation a ring of tight junctions additionally forms around the adherens junctions between the MC and the oocyte surface.

Adherens and tight junctions associate with the actomyosin cytoskeleton (reviewed in Arnold et al., 2017; Quiros and Nusrat, 2014). As described above, actin was strongly enriched at the MC-oocyte contact point from early stage III onward (Fig. 6A) and this thick domain largely overlapped with β-catenin proteins (Fig. 6H’). To determine how non-muscle myosin was distributed relative to the actin cytoskeleton in the early oocyte and the MC, we used the *Tg(actbl:myll2.l-eGFP)* transgenic line that expresses the light chain of non-muscle myosin II fused to GFP (Maître et al., 2012). Non-muscle myosin II organized as a thin layer all around the oocyte cortex together with F-actin in early stage III oocytes (see Fig. 7A for Myl12.1-gfp and Fig. 6A for F-actin). Intriguingly, as the oocyte development proceeded and the MC started to grow and differentiate, a domain free of non-muscle-myosin II was visualized corresponding to MC-oocyte contact zone (Fig. 7B, white bracket). This gap was specific to the non-muscle myosin II staining since the actin cytoskeleton organization appeared uninterrupted (see Fig. 6A, B, H, J, K’). The size of the non-muscle myosin II-free zone expanded as the MC became bigger (Fig. 7C, white bracket). In the MC itself, the activated, phosphorylated form of non-muscle myosin II (pNMII) localized at the membrane from early stage III oocyte onwards (Fig. 7D). At mid-stage III, when the MC extends drastically in length, activated nonmuscle myosin II co-localized with F-actin at the lateral membrane of the MC (Fig. 7E, E’). At late stage III, however, once the MC was close to reach its final size, both F-actin and phospho-myosin were barely detectable at the membrane (Fig. 7F, F’), similarly to β-catenin and E-cadherin (Fig. 6I’, K’). This dynamic spatio-temporal distribution of non-muscle myosin II in the oocyte and in the MC suggest that actomyosin-mediated tension might play an important role in the specification of the MC precursor and in the activation of Taz.

**Figure 7.**
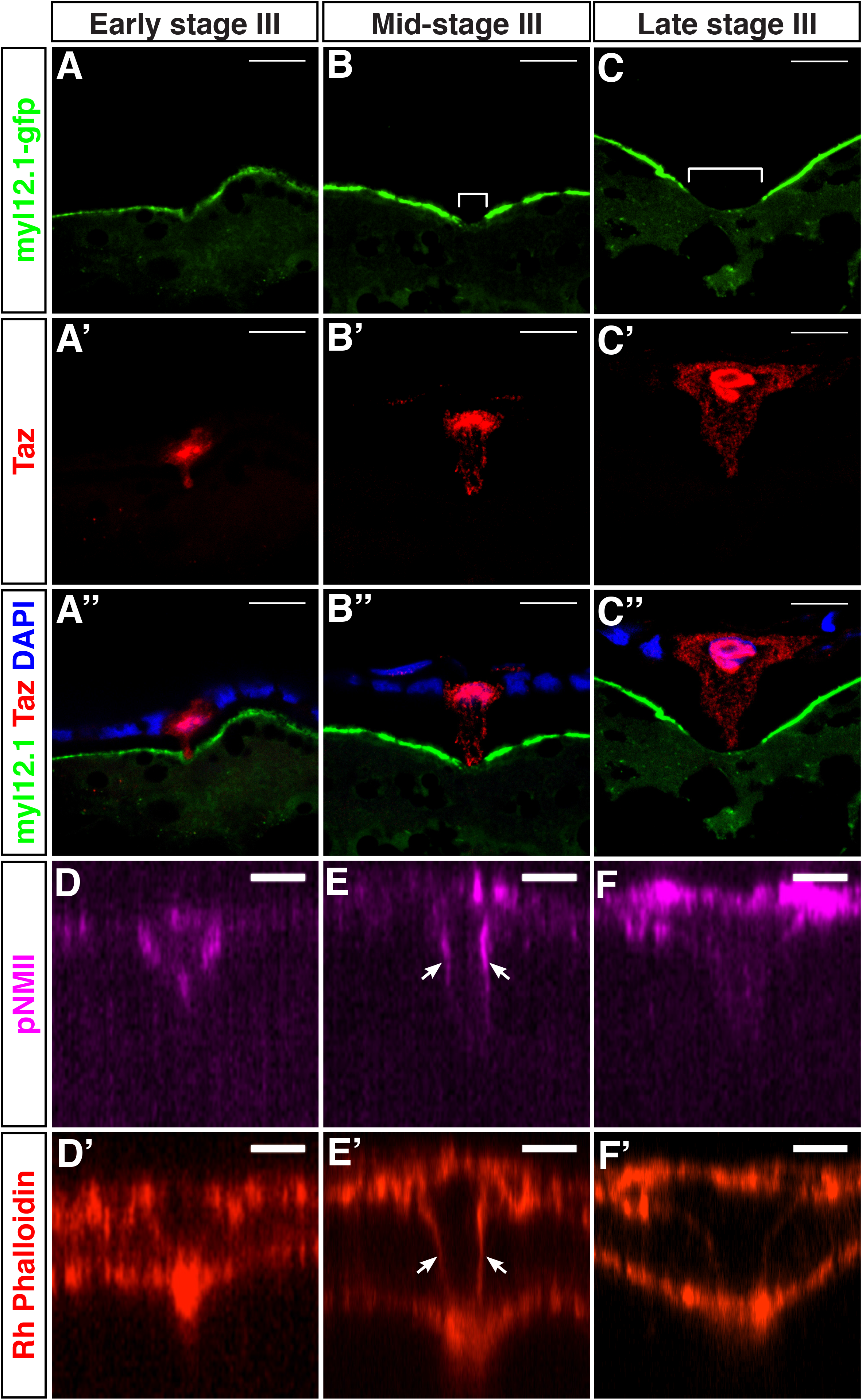
Non-Muscle Myosin localization in the oocyte during MC morphogenesis. (A-C’’) Confocal images of cryosectioned ovaries displaying the temporal changes in the localization of Myl12.1-GFP localization at the animal cortex of the oocyte (N=2, n=34). During stage III, a wider zone with strongly reduced Myl12.1-GFP forms (white bracket in A-C). Taz is used as a MC marker (A’-C’). DAPI is used to label the FCs and MC nuclei as shown in the merged images (A’’-C’’). (D-F’) Whole mount immunofluorescent images stained with an anti-phosphorylated non-muscle myosin II antibody (pNMII, magenta, n=27) at the indicated stages of oogenesis. Solid arrow in the panel E and E’ indicates the localization of the pNMII and actin at the lateral membranes of the MC, respectively. Rhodamine-Phalloidin is used as to independently mark the MC membranes (red, D’-F’). Scale bar: 10 μm in A-F’.

Ultrastructural studies in Medaka and in the Chum Salmon have shown that the MC is particularly enriched in desmosome-associated intermediate filaments (Kobayashi and Yamamoto, 1985; Nakashima and Iwamatsu, 1989). To determine whether this is also the case in zebrafish, we stained stage III follicles for the desmosomal marker Desmoplakin (Dsp) and the intermediate filament component Keratin 18 (Krt18) (Fig. 8). Intriguingly, both Desmoplakin and Keratin 18 were strongly enriched in the MC in all analyzed oogenesis stages when compared to the rest of the FCs (Fig. 8 A-J). Desmoplakin was present as aggreagates in the entire MC cytoplasm and was enriched at the base of the MC cytoplasmic process at the contact point with the oocyte (Arrow in Fig. 8 A-D). This localization was similar to β-catenin and E-cadherin localization (Fig. 6 G, I, K). In contrast, Krt18 organized as a thick coat around the nucleus in the MC cytoplasmic process at all stages of morphogenesis (Fig. 8E-J). Krt18 staining was always weaker in the MC cytoplasmic process but this does not exclude the presence of other Keratins (Fig. 8G, G’). This result suggests that intermediate filaments play an important functional and structural role to mechanically support the MC body during its elongation.

**Figure 8.**
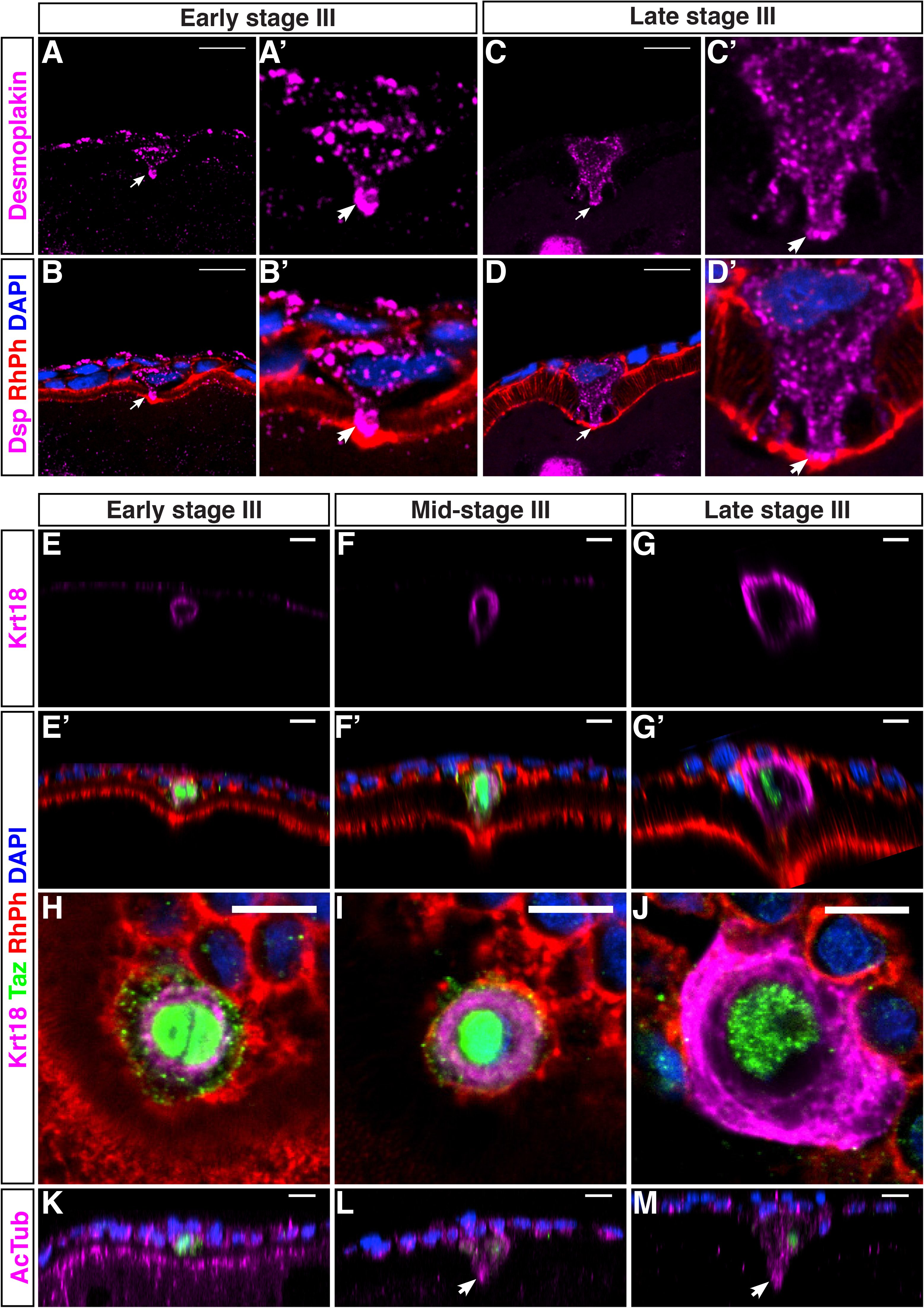
Desmoplakin, Keratin 18 and Acetylated-tubulin localization pattern during morphogenesis of the MC. (A-D’) Confocal images of cryosectioned ovaries stained with an anti-desmoplakin antibody (magenta), Rhodamine-Phalloidin (F-actin, red) and DAPI (nuclei, blue) (n=13) showing that desmoplakin is specifically enriched in the MC from early stages of its morphogenesis and is present at the contact point between the MC and the oocyte. It also colocalizes with F-actin at the base of the MC (solid white arrow). (E-J) Whole mount immunofluorescence with an antibody against the intermediate filament protein Krt18 (n=11). (E-G’) are orthogonal XZ views of confocal stacks shown in (H-J). Like Desmoplakin, Krt18 is also enriched specifically in the MC from early stage III onwards (E-G), forms a thick layer around the nucleus, but is absent in the MC cytoplasmic process. The Taz antibody (green), DAPI (blue) and Rhodamine-Phalloidin (red) were used to counterstain the MC, the nuclei and the membranes, respectively. (K-M) XZ sections of follicles at the indicated stages stained in wholemount with an anti-acetylated tubulin antibody showing the localization pattern of stable microtubules in the MC (n=16). Stable microtubules are specifically enriched at the base of the cytoplasmic process where the MC contacts the oocyte surface from mid-stage III onward, (arrowhead in L and M). Scale bar: 10 μm in AM.

Since Krt18-containing intermediate filaments were concentrated in the cell body of the MC, we tested whether microtubules might play a structural role, during the growth and maintenance of the MC process, in pushing down against the oolemma, as shown in Medaka and the Chum salmon oocytes (Kobayashi and Yamamoto, 1985; Nakashima and Iwamatsu, 1989). For this purpose, we stained whole-mount follicles during MC morphogenesis with an antibody against acetylated Tubulin to label stable microtubules. From mid-stage III onward, stable microtubules were enriched at the tip of the MC cytoplasmic process (white arrow in Fig. 8L, M). This suggests that stable microtubules might be important for the growth of the MC process in the oocyte cortex and its resistance to extrinsic mechanical stress.

In summary, we have generated the first phenotypic map of the MC morphogenesis at the molecular level. This characterization of the MC show that it is anchored to the oolemma throughout morphogenesis via β-catenin and Desmoplakin-containing junctions, associated with actin filaments (Fig. 9A-F). In addition, tight junctions reinforce the MC-oocyte link (Fig. 9E-F). Our data further reveal that actomyosin (Fig. 9D-F), microtubules (Fig. 9B-C) and intermediate filaments (Fig. 9A-C) likely cooperate downstream of Taz function to: (i) actively drive the drastic changes in cell shape and size, (ii) stabilize these changes and (iii) provide the MC mechanical resistance to pressure originated from the forming vitelline membrane, the neighboring FCs and the oocyte.

**Figure 9.**
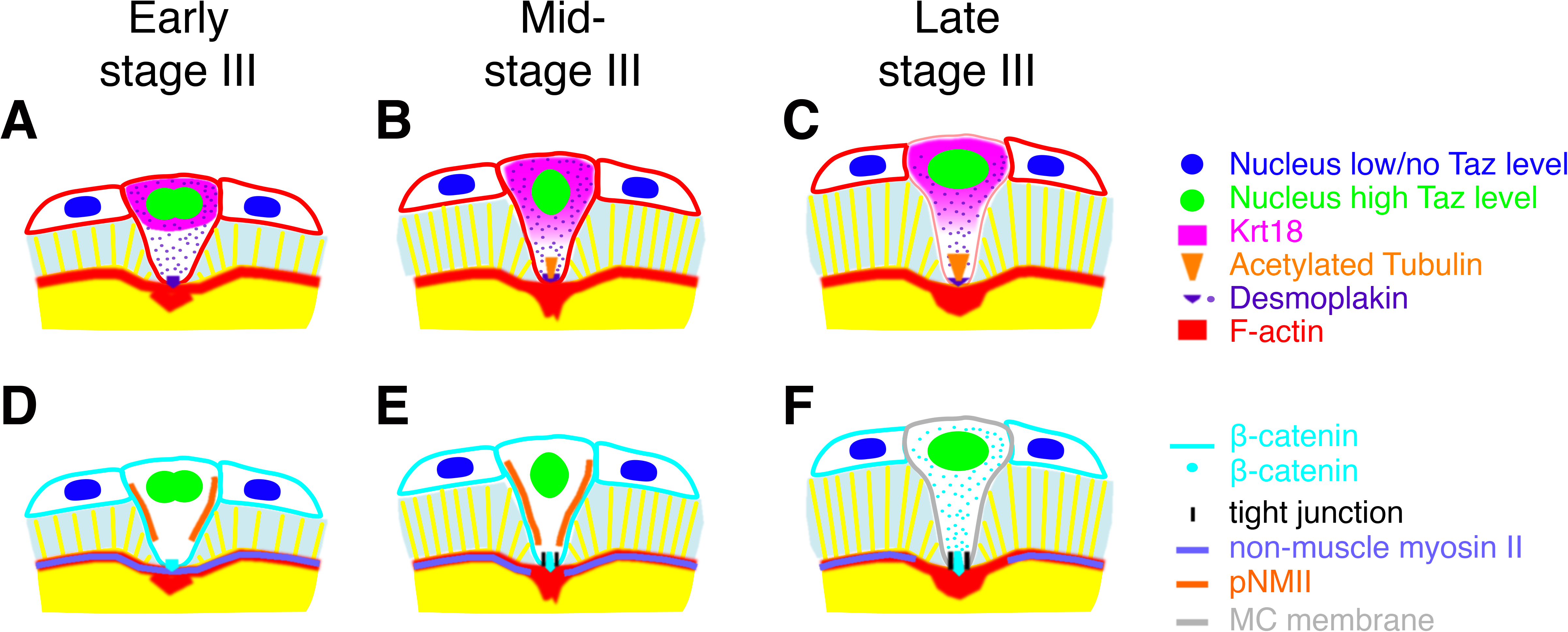
Schematic molecular map summarizing the dynamic localization of different molecular markers in the MC during stage III of oogenesis. (A-F) summarize the localization of the different molecular markers looked at in this study. The markers have been separated into two sets (A-C and D-F) as indicated on the right side to increase the clarity. The long stage III has been subdivided into three periods for a better temporal resolution of the MC morphogenetic process: early (A, D), mid - (B, E) and late (C, F) stage III.

## Discussion

In this study, we show that female lacking the Hippo pathway effector Wwtr1/Taz are infertile because their eggs lack a micropyle and thus cannot be fertilized by a sperm. We further show that Taz is required for the specification of the MC among all FCs surrounding the oocyte and constitutes the first MC biological marker described to date. Finally we perform the first molecular characterization of the MC in zebrafish by analyzing multiple cytoskeletal and junctional markers. This allowed us to identify two additional players, in addition to Taz, that specifically differentiate the MC from other FCs: the intermediate filament component Keratin 18 and the desmosomal protein Desmoplakin. Altogether these results lead us to propose a hypothetical model on the upstream mechanism that specifically activates Taz in one particular cell (Fig. 10, see below), and the mechanism downstream of Taz activity that leads to the differentiation of the MC (Fig. 9). Here, we provide insights into the genetic basis of micropyle biogenesis. It will also help to explore the evolutionary fate of somatic factors during development and reproductive strategies selected for in animals, as well as, for functional interrogation of novel regulators playing a role during this process.

**Figure 10.**
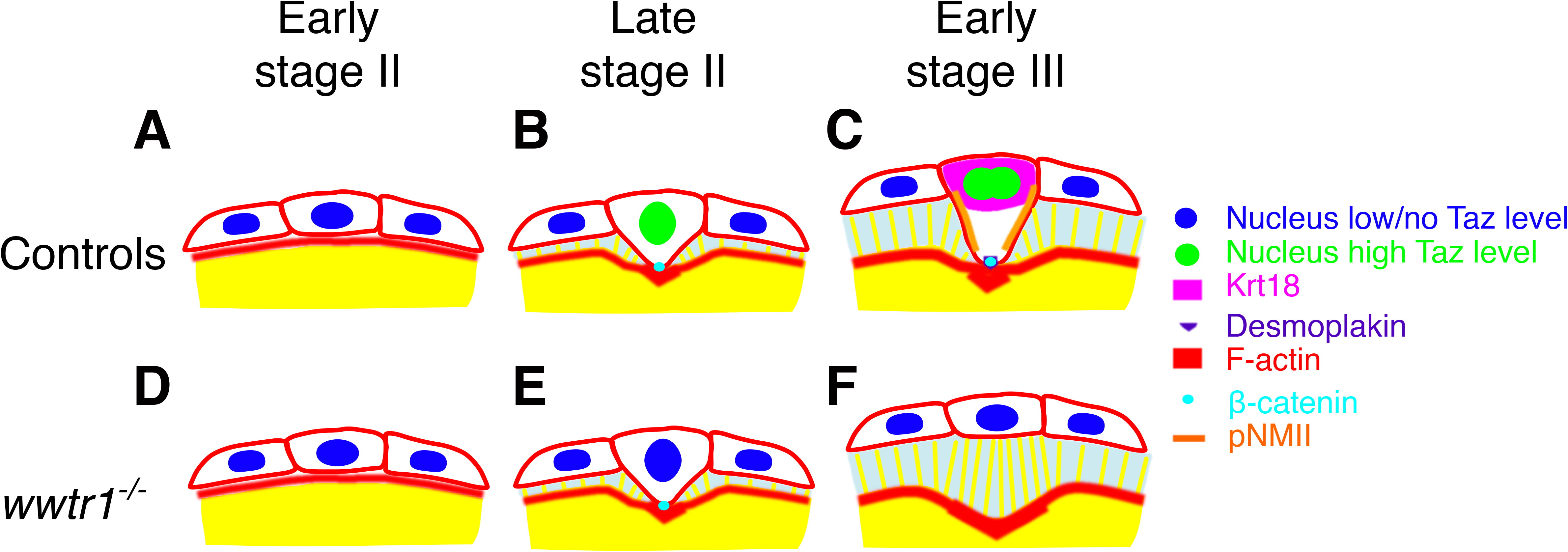
Working model of the micropylar cell fate specification. Working model based on the results obtained in the present study. Our hypothetical model described the specification of a single MC at the animal pole by Taz functioning downstream of physical cues originated in the oocyte in WT (A-C) and its failure in *wwtr1^-/-^* follicles (D-E).

### Taz functions as a major regulator of the micropylar cell fate

We show that Taz activity: (i) is specifically upregulated in the future MC precursor during early oogenesis, before its obvious and dramatic morphological changes, and (ii) is required for MC differentiation and micropyle formation. These findings indicate that Taz, as a transcriptional co-activator, is a major regulator of this specific somatic cell type. But is Taz activity sufficient to make any FC competent to adopt a MC fate? Unfortunately, expressing *wwtr1* mosaically and specifically in the FC layer during oogenesis is experimentally very challenging because of the absence of a FC-specific promoter. Interestingly, however, we show here that *buc* mutant oocytes represent a valuable and experimental model to address this question (Fig. 5F, G). In *buc* mutant oocytes, polarity fails to be established and multiple micropyles form (Marlow and Mullins, 2008). We show here that this phenotype correlates with multiple and ectopic Taz expressing MCs. The study of the tight communication established between the oocyte and FCs during the zebrafish folliculogenesis have shed lights into the underlying mechanism regulating oocyte development progression and survival (Li and Ge, 2011; Yan et al., 2017). However, the identification of reciprocal signaling exchange governing micropyle biogenesis has not been reported yet. Our results strongly support an early role for Taz in the cascade leading to the differentiation of the MC, and makes this factor an excellent candidate to translate the spatially restricted information previously defined in the oocyte into positional information in the FC layer. To understand how Hippo signaling components participate in the dual cross-talk between maternally loaded factors into the oocyte and the MC precursor will require further investigations.

### The MC is not required for oocyte polarity maintenance

In *buc* mutant early oocytes, mRNAs that normally localize specifically at the vegetal (*dazl*) or at the animal (*pou2*) pole fail to do so, being either diffusely or radially distributed, respectively. In the mutant oocyte the *vg1* mRNA, which normally localizes at the animal pole, localizes in discrete peripheral cytoplasmic domains in correlation with each ectopic micropyle. This led to the hypothesis that the anchoring of the *vg1* transcript to the animal pole, might be influenced by the MC via its interaction with the oolemma (Marlow and Mullins, 2008). Surprisingly, the *vg1* mRNA animal localization is unmodified in the *wwtr1* mutant oocytes (Supp Fig. 2) indicating that the MC emergence is not required to localize this transcript. We examined several oocyte polarity markers, including *cyclinb1, dazl* or Zorba:GFP, however, their spatial localization was similarly unaffected in *wwtr1^-/-^* oocytes. In conclusion, oocyte polarity establishment limits the number of MCs to one, but MC differentiation is not required to maintain the AV coordinates in the developing oocyte. Future phenotypic analysis and functional studies will decipher the potential interplay between the animal cytoplasmic domain and Taz during the FC to MC transition.

### What is the nature of the spatiotemporal signals required to specify a unique MC in the early oocyte?

The MC is an ideal model to address the long-standing question of what makes one cell in an initially homogeneous cell layer become different from its neighbors in a particular developmental stage. The asymmetry of the oocyte plays a crucial role to both localize the MC at the animal pole and prevent the formation of ectopic MCs (Marlow and Mullins, 2008). How this polarity information translates into the specification of one cell in the overlying FC layer is still unknown. Our results show that nuclear translocation and enrichment of Taz in one of many cells within the FC layer leads to MC formation. Conversely, in *wwtr1*/Taz loss-of-function oocytes, the MC does not form. It makes the *wwtr1* mutant a suitable and attractive model to explore the genetic underpinnings behind the FC to MC transition.

The nature of the signal from the oocyte to the follicular cell layer is not clear but a close-range signal or even direct cell contact, rather than a broadly diffusible signal, has been proposed (Heim et al., 2014). Taz, and its paralog Yap1, are known to respond in many different contexts to changes in cell adhesion, actin cytoskeleton reorganization or mechanical forces by translocating into the nucleus (reviewed in Low et al., 2014; Sun and Irvine, 2016; Varelas, 2014; Yu and Guan, 2013). It is therefore tempting to speculate that the increased Taz activity we observed during oogenesis in the future MC might be induced by a mechanical signal originated in the oocyte. At the onset of oogenesis, the follicular cells are in direct contact with the oocyte (Fig. 5J and 10A, B). As the oocyte grows, the microvilli at its surface increase in length, and vitelline membrane material accumulates in between. Thus, the distance between the oocyte and the FC layer also increases (Fig. 5M and 10B, E). At the animal pole however, the MC precursor remains in contact with the oocyte and thus elongates to accommodate the increased distance between the FC layer (to which it belongs) and the oocyte (to which it is attached) (Fig. 5M and 10B, E). We propose that either the adhesion to the oocyte or changes in cellular morphology could induce the nuclear translocation of Taz in the MC precursor as previously in other cell types (Fig. 10B) (Aragona et al., 2013; Dupont et al., 2011; Wada et al., 2011; Zhao et al., 2007).

We also report the formation of an indentation at the animal pole of the oocyte beneath the MC (Fig. 4A, K). While this indentation could be caused by the MC pushing forces against the oolemma, we observed an even more pronounced, and earlier indentation in *wwtr1^-/-^* oocytes (Fig. 4B, S). This suggests the existence of forces in the oocyte cortex that would pull on the facing FC. These pulling forces combined to the attachment of the referred FC to the oocyte would lead to the enrichment and nuclear translocation of Taz in this selected cell. The presence of a cortical actomyosin network surrounding the oocyte beneath the oolemma supports the idea of a contractile cortex capable of generating forces.

The nuclear translocation of Yap/Taz upon cell stretching or cell shape changes has been shown to depend on an increase in actomyosin-mediated cellular tension (Aragona et al., 2013; Dupont et al., 2011; Wada et al., 2011). Accordingly we observed phospho-myosin localized at the membrane of the MC during early differentiation stages (Fig. 7D-E, Fig. 10C), while this membrane localization decreased at later stages (Fig. 7F). We thus propose that the attachment of the future MC to the oocyte animal pole, in combination to pulling forces from the oocyte, play an important role in regulating Taz protein levels and nuclear translocation. Taz would then induce the expression of genes required for the growth of the MC (see below). Thus, transcription and translation of Taz targets would enable the MC to accommodate its shape to global morphological changes between the oocyte and the FC, and reduce mechanical tension. Genome-wide expression profiles studies in temporal scales emerge as a promising approach to support this hypothesis. It will also set the basis for future interrogation of their contribution, upstream or downstream of Taz, to MC differentiation and maintenance for successful fertilization.

In *wwtr1^-/-^* oocytes, the pulling forces would remain but Taz would not be activated and therefore the MC would not be specified (compare Fig. 10E to Fig. 10B). Since Taz activation is likely promoting a reinforcement of the MC-oocyte attachment (Fig. 5G-K), this would limit the indentation depth at the animal pole in the wild-type oocyte. We propose that this reinforcement, downstream of Taz function, fails to be executed in *wwtr1^-/-^* follicles. As a consequence, the FC layer fully detaches from the oolemma at the animal pole. This lack of attachment combined to pulling forces from the oocyte would lead to the oocyte cortex falling back and to the pronounced indentation observed in *wwtr1^-/-^* oocytes (Fig. 10 C, F). Altogether our observations support the idea of a mechanical link between the oocyte and the MC and suggest that the signal from the oocyte leading to the specification of one MC could be mechanical.

Intriguingly, we observed a delocalization of the cortical non-muscle myosin at the animal pole indentation just beneath the differentiating MC in WT oocytes (Fig. 7A-C). The cortical myosin layer was still intact in early stage III oocytes when the MC was easily distinguishable from the rest of the FCs. This strongly suggests that this delocalization is initiated after the MC has started to enlarge, and could be a consequence of the MC process pushing on the oolemma. Alternatively, the strong reduction in cortical non-muscle myosin at the animal pole could be intrinsically triggered by the oocyte to modify the physical properties of the cortex specifically at the MC-oocyte contact site. A reduction of non-muscle myosin would indeed reduce cortical tension and soften the oocyte cortex in this area, thus facilitating the pushing of the MC process into the oolemma. Such a delocalization of cortical non-muscle myosin has been shown to decrease cortical tension in mouse oocytes, a prerequisite to properly position the meiotic spindle (Chaigne et al., 2015). Interestingly, nonmuscle myosin exclusion in mouse oocytes correlates with a thickening of cortical F-actin, as we found at the animal pole of the zebrafish oocyte. The cortical actin thickening in mouse oocytes has been shown to be nucleated by the Arp2/3 complex and to trigger the delocalization of non-muscle myosin from the cortex (Chaigne et al., 2015). Whether the delocalization of non-muscle myosin from the zebrafish oocyte cortex is triggered by the actin thickening, is linked to oocyte polarity or is initiated by the MC morphogenesis will require further analysis.

### What are the events downstream of Taz driving differentiation of the MC?

One question that our work raises is which cellular events are activated downstream of Taz, that make a FC differentiates into the very specialized MC. There are two major differences between these two related cells: their size and their shape. Concerning the increase in size, two factors have been mainly implicated in cell growth regulation: the transcription factor Myc and the mTOR signaling pathway (reviewed in Lloyd, 2013). Interestingly, Myc has been shown in *Drosophila* to function as a critical effector of cellular growth downstream of Yki, the homolog of Yap/Taz (Neto-Silva et al., 2010). Similarly, several studies have shown that the Hippo-YAP pathway acts as an upstream regulator of mTOR and that these two pathways coordinate to control cell size and growth (James et al., 2009; Liu et al., 2017; López-Lago et al., 2009; Santinon et al., 2016; Tumaneng et al., 2012). It will therefore be interesting to test if the strong increase in size of the MC could be due to an increase in Myc or mTOR activity downstream of Taz. Interestingly, Keratin cytoskeletal proteins, including Krt18, have been shown to regulate protein synthesis and cell growth upstream of the Akt/mTOR signaling pathway in keratinocytes (Kim et al., 2006). Since we show here that Krt18 is specifically and highly enriched in the MC specifically, Krt18 might be involved in the strong size increase of the MC.

In addition to the obvious increase in size, the MC acquires a very characteristic mushroom-shape during its morphogenesis. Cell shape changes are often associated with dynamic properties of the actomyosin cytoskeleton (reviewed in Murrell et al., 2015; Rauzi and Lenne, 2011; Salbreux et al., 2012), and we show here that both actin and myosin are dynamically regulated during MC morphogenesis. Importantly, the MC achieves its peculiar shape over the period of a few weeks. This is in strong contrast with cell shape changes during developmental processes that occur over minutes or hours. This raises the question of whether intrinsic cellular mechanisms take place in the MC to shape and maintain the desirable morphology over such a long period of time. In addition to the intermediate filament component Krt18, we show here that the desmosomal protein Desmoplakin is also highly enriched in the MC as compared to other FCs (Fig. 8A-D). These results suggest that intermediate filaments anchored to desmosomes might play a role in the acquisition and maintenance of the MC final shape. Krt18 forms a rim around the MC nucleus that thickens as the MC grows in size (Fig. 8E-J). We propose that intermediate filaments provide the MC mechanical strength to support the plasma membrane to withstand the pressure from the forming vitelline membrane around over longer time period. This is in agreement with the structural role of intermediate filaments in providing mechanical strength, including plasma membrane support to cells (reviewed in Loschke et al., 2015; Quinlan et al., 2017). In addition, we observe enrichment in acetylated tubulin in the MC cytoplasmic process that pushes on the oolemma (Fig. 8K-M). Acetylation of microtubules has recently been proposed to enable microtubules to better resist mechanical stress, in particular over long period of time and during cell ageing (Janke and Montagnac, 2017; Portran et al., 2017). Thus, changes associated with MC differentiation are compatible with a slow developmental process and with the requirements of the MC to withstand and mechanically modify the vitelline membrane around it.

## Conclusions

We have identified a novel, unique and essential role for the Hippo pathway effector Taz in the specification of the MC fate. In the absence of Taz activity, a MC fails to differentiate within the FC layer. As a consequence, the chorion of mature eggs lack a micropyle leading to female infertility. We identify Taz as the first *bona fide* marker of the MC identity. The strong enrichment of Taz protein in the future MC before obvious morphological changes strongly suggests that Taz is a major regulator of the MC fate in zebrafish. Furthermore, based on our molecular characterization of the MC and its contact point with the oocyte, we propose the following model (Fig. 10): (i) the oocyte AV polarity generates differences at the animal pole cortex that translate into an increased adhesion with the facing FC, (ii) as the oocyte grows and microvilli at its surface elongate, all FCs move away from the oocyte surface except the cell at the animal pole which, as a consequence, elongates. (iii) The attachment to the oocyte and/or the elongation lead to an increase in level and nuclear translocation of Taz in a single cell that becomes the MC precursor. Downstream of Taz (Fig. 9), the MC differentiation program is induced. This includes: (a) reinforcement and stabilization of the anchoring of the MC to the oolemma via adherens junctions and desmosomes, (b) formation of an intermediate filament coat that gives the cell mechanical resistance and (c) localization of stable microtubules that are likely to be important for the MC cytoplasmic process to push on the oocyte plasma membrane. Future studies will be necessary to identify the precise molecular mechanisms that are activated upstream and downstream of Taz that control the specification and differentiation of the MC, respectively.

## Material and Methods

### Zebrafish husbandry

Procedures involving animals were conducted according to institutional guidelines and approved by the German authorities (veterinary department of the Regional Board of Darmstadt). Adult zebrafish were maintained in a constant circulating system with a 14 hour light/10 hour dark cycle. The previously characterized mutant *taz^mw49^* (Miesfeld et al., 2015), and transgenic *Tg(4xGTIIC-d2EGFP)^mw50^*(Miesfeld and Link, 2014) and *Tg(actb2:myl12.1-eGFP)^e2212Tg^* (Maître et al., 2012) lines were used in this study. *In vitro* fertilization was performed according to standard procedures (Westerfield, 2007)

### Generation of mutant and transgenic zebrafish lines

The mutant line *wwtr1^fu55^* was generated using TALEN-induced mutagenesis strategy. A target site in the first exon and the corresponding left and right TALENs were designed using the online software MojoHand (http://www.talendesign.org). The TALENs were cloned using the TALEN repeat array plasmid library (Addgene kit #1000000024) and Golden Gate Assembly (Cermak et al., 2011). The detailed protocol is available on the Addgene website (https://www.addgene.org/static/cms/filer_public/98/5a/985a6117-7490-4001-8f6a-24b2cf7b005b/golden_gate_talen_assembly_v7.pdf). The plasmids containing the RVDs fused to Fok1 were linearized with BamHI and corresponding RNAs were *in vitro* transcribed using the T3 mMessage mMachine Kit (Ambion by Life Technologies GmbH, Darmstadt, Germany). The left and right TALEN mRNAs were co-injected in one-cell stage embryos and several mutant alleles were recovered by amplifying a region encompassing the target site by PCR using the primers *wwtr1-tall-e1-fow* and *wwtr1-tall-e1-rev* (see supp. Table) and digesting it with BspHI which is present in the spacer in the *wwtr1* WT sequence. Zygotic mutants obtained from an incross of heterozygous mutant fish were raised to adulthood to obtain F3 maternal-zygotic (MZ) mutants. Zygotic WT “cousins” obtained from the incross of the heterozygous mutant fish were raised and used as littermate controls for the MZ mutant fish.

The *zp0.5:eGFP-zorba* transgene was generated by amplifying the *-0.5zp3b* promoter from a *-0.5zp3b:GFP* construct (Onichtchouk et al., 2003) and cloning it into a vector containing recognition sites for the Tol2 transposase. Full-length *zorba* cDNA was amplified from total zebrafish cDNA and cloned into the vector containing the *-0.5zp3b* promoter and the Tol2 sites. Positive oocytes expressing the eGFP-Zorba fusion protein were isolated from founder F0 females but positive F1 could not be recovered. The construct was injected at a concentration of 40 ng/μl along with the Transposase mRNA (50 ng/μl). Transposase mRNA was synthesized with the mMessage mMachine Sp6 Polymerase Kit (Ambion) from pCS2FA-transposase (Kwan et al., 2007).

### Mature Egg activation and in vitro fertilization

Mature eggs were obtained by gently squeezing the belly of gravid females and were activated in E3 medium. The micropyle was then either observed directly or stained with Coomassie Brilliant Blue (Yanagimachi et al., 2013). *In vitro* fertilization was performed according to standard protocols (Westerfield, 2007).

### Ovary dissection and immunostaining on cryosections

Adult zebrafish were euthanized using an overdose of tricaine according to the institutional guidelines approved by the German authorities. To dissect the ovary out, freshly euthanized females were dissected using a sharp pair of forceps. The ovarian tissue was quickly rinsed in L-15 Medium (pH=9) and then used for further processing.

For cryosections, dissected ovaries were fixed with 4% PFA in PBS (pH 7.4) overnight at 4°C and washed with PBS five times for 5 mins each. Whole ovaries were then transferred to the following solutions and allowed to equilibrate for one day: 25% sucrose in PBS, 35% sucrose in PBS, 1:1 O.C.T. (TissueTek): 35% sucrose in PBS and 100% O.C.T. The ovaries were then transferred to a mold containing fresh O.C.T. and frozen on a metal block that was stored at –80°C. 10 micron-cryosections were obtained using a cryotome, with the block at –15°C and the chamber at –20°C, were placed on Superfrost UltraPlus slides (Thermo Scientific, J3800AMNZ), and air-dried at room temperature for at least 2 hours and stored at –80°C.

### Oocytes isolation and fixation for whole-mount immunostaining and In Situ Hybridization

Dissected ovaries were digested with Collagenase (0.25 mg/mL) for 10 minutes and individual ovarian follicles were gently dissociated using a cut 1 mL tip. Dissociated follicles were washed with L-15 medium (pH 9) 3-4 times. L-15 medium was removed completely and follicles were acid-fixed overnight at 4°C or for 2 hours at RT with 2 mL of 5% Formaldehyde in PBS and 8-10 drops of Glacial Acetic Acid (1 drop is approximately 50 μl, final concentration of the acid - 5% - 8%) (Fernández and Fuentes, 2013). Fixed follicles were washed 5 times with PBS containing 0.1% Tween-20 (PBST). In Fig. 6 (pNMII/phalloidin staining), ovarian follicles were fixed in 4% PFA in PBS overnight at 4°C.

Acid-fixed or PFA-fixed follicles were blocked with PBDT-I (1X PBS, 1% BSA, 1% DMSO and 0.5% TritonX-100) containing 2% Normal Goat Serume (NGS) for at least 30 min. Follicles were incubated in primary and secondary antibodies either overnight at 4°C or 4 hours at RT. After antibody incubation, the follicles were washed 5 times for 20 minutes in PBDT-I. The following antibodies were used: mouse anti-β-catenin (1:200, Sigma C7207), rabbit anti-Taz (1:200, Cell Signaling, D24E4) (Miesfeld et al., 2015), mouse anti-acetylated tubulin (1:200, Sigma T6793), mouse anti-Desmoplakin (ready-to-use, Progen, 651155), mouse anti-Krt18 (1:200, Thermo, MA1-06326), rabbit anti-pNMII (1:50, Cell Signalling 3671S), mouse anti-GFP (1:500, Clontech 632381), rabbit anti-GFP (1:500; Torrey Pines Biolabs), mouse anti-E-Cad (1:200, BD Sciences, 610182) and mouse anti-ZO1 (1:500; Invitrogen). Alexa dye-conjugated antibodies (Thermo Fischer Scientific) were used at 1:500 dilution. Immunofluorescence on cryosections were performed according to the protocol published on Abcam’s website (http://www.abcam.com/protocols/immunostaining-paraffin-frozen-free-floating-protocol). Rhodamine-Phalloidin (Molecular probes R415) was used at 1:100 dilution. The sections were blocked in 5% NGS in PBDT-II (1X PBS, 1% BSA, 1% DMSO and 0.025% TritonX-100) and all the washes were performed with PBDT-II.

As many antibodies do not work after ISH, this procedure were performed after immunostaining on acid-fixed ovarian follicles whenever necessary (Elkouby and Mullins, 2016). Immunostained follicles were post-fixed in 4% PFA in PBS for 20 minutes at RT followed by 5 washes of 5 min in PBST. ISH were then performed according to standard protocol starting at the pre-hybridization stage (Thisse and Thisse, 2008). Hybridization was performed at 60°C overnight. The following probes and antibodies were used: *cyclinb1* (1:200) (Kondo et al., 1997) to mark the animal pole, *dazl* (1:200) to mark the vegetal pole and the Balbiani Body (Kosaka et al., 2007), *vg1* (1:200) to mark the animal pole just beneath the MC. All the probes used in this study are DIG-labeled (DIG RNA Labeling Mix, Roche, 11277073910). Anti-DIG-AP (Roche, 11093274910) antibody was used and FastRed (Roche, 11496549001) was used as substrate in 0.1 M Tris buffer with 0.1% Tween-20 (pH 8.2) to detect *cyclinb1* and *dazl* transcripts. For *vg1* transcripts, NBT (Fermentas, R0482) - BCIP (Fermentas, R0822) was used in NTMT buffer (100mM NaCl, 50mM MgCl_2_, 100mM Tris-Cl, 0.1% Tween-20, pH 9).

### Microscopy and image processing

Ovarian follicles were staged according to Selman et al., 1993. Individual follicles were mounted on 24×60 mm coverslips in a 0.8% low-melting agarose drop diluted in E3 medium. Imaging was peformed on a Nikon W1 Spinning Disc Microscope with the following objectives: 10X (NA 0,45, WD 4mm) air objective or 20X (NA 0,95, WD 0,95mm), 40X (NA 1,15, WD 0,60mm) and 60X (NA 1,20, WD 0,30mm) water objectives. Images were processed using the FIJI software.

### RNA Extraction, cDNA synthesis and RT - PCR

Total RNA was extracted with Trizol from samples collected at the following developmental stages: mature oocyte, 256-cell stage, sphere, 60% epiboly and 28 hpf. 1 μg of total RNA was reverse-transcribed using the Superscript III (Invitrogen). Primers sequences for RT-PCR are shown in the Supp. Table 1.

## Acknowledgment

We are grateful to Sabine Götter in Prof. Wolfgang Driever’s lab (Freiburg), Denise Werner and Tore Dittrich (Frankfurt) for excellent fish care. We thank Daria Onichtchouk and Roland Dosch for valuable input at the beginning of the project, and Prof. Stefan Eimer for critical reading of the manuscript.

## Author Contributions

C. D. and V.L. designed the experiments and analyzed the data. CD, AN, PK and SG performed the experiments. R.F. and M.C.M. participated in experimental design, data analysis and provided reagents. C.D. and V.L. prepared the figures and wrote the manuscript and all authors edited the paper prior to submission. V.L. supervised and coordinated the project.

